# Sequencing whole genomes of the West Javanese population in Indonesia reveals novel variants and improves imputation accuracy

**DOI:** 10.1101/2024.06.14.598981

**Authors:** Edwin Ardiansyah, Anca-Lelia Riza, Sofiati Dian, Ahmad Rizal Ganiem, Bachti Alisjahbana, Todia P Setiabudiawan, Arjan van Laarhoven, Reinout van Crevel, Vinod Kumar, ULTIMATE consortium

## Abstract

Existing genotype imputation reference panels are mainly derived from European populations, limiting their accuracy in non-European populations. To improve imputation accuracy for Indonesians, the world’s fourth most populous country, we combined Whole Genome Sequencing (WGS) data from 227 West Javanese individuals with East Asian data from the 1000 Genomes Project. This created three reference panels: EAS 1KGP3 (EASp), Indonesian (INDp), and a combined panel (EASp+INDp). We also used ten West-Javanese samples with WGS and SNP-typing data for benchmarking. We identified 1.8 million novel single nucleotide variants (SNVs) in the West Javanese population, which, while similar to the East Asians, are distinct from the Central Indonesian Flores population. Adding INDp to the EASp reference panel improved imputation accuracy (R2) from 0.85 to 0.90, and concordance from 87.88% to 91.13%. These findings underscore the importance of including Indonesian genetic data in reference panels, advocating for broader WGS of diverse Indonesian populations to enhance genomic studies.

## Introduction

Genome-wide association studies (GWAS) have played a pivotal role in advancing our understanding of the genetic underpinnings of diseases over the past decade^1,2^. Genotype imputation, has gained paramount importance in GWAS, by inferring unobserved genotypes^3^. To facilitate imputation, the 1000 Genomes Project (1KGP3) has furnished the genome sequencing data required for constructing reference panels^4^. However, the current repository of genotypes is dominated by genotype information from the European populations, thereby inadequately representing global diversity. Consequently, imputing genotypes for individuals from non-European populations can result in reduced accuracy due to differences in genetic variation between populations^5^. Prior research has also indicated significant disparities in imputation performance when employing common reference panels across diverse populations^6,7^. Such inaccuracies particularly limit the possibility of detecting genetic association at low-frequency variants, leading to incomplete understanding of the genetic architecture underlying complex diseases in ethnically diverse populations.

Indonesia, the world’s fourth most populous country, with over 17,000 islands and a diverse population^8^ is currently underrepresented in human genomic studies. The closest super-population in the 1KGP3^4^ database to Indonesia is the East Asian (EAS) panel, composed of populations from China, Japan, and Vietnam. However, this East Asian panel may inadequately represent the genetic landscape specific to Indonesia. Addressing this shortfall, the GenomeAsia 100K (GAsP) consortium has sequenced 1,739 individuals from Asia^9^. GAsP included 68 individuals from the central parts of Indonesia, but does not capture the rich genetic diversity present throughout the archipelago.

In the present study, we examined the largest cohort of genomes from individuals originating from West Java, the most populous province of Indonesia with 50 million inhabitants. Our primary objective was to identify novel genetic variants specific to this region. Furthermore, we investigated the added value of incorporating our WGS data in the current Asian reference panels to improve the accuracy of imputation.

## Results

### Whole genome sequencing of West Java population identifies 1.8 million novel SNVs

We performed whole genome sequencing (WGS) on 239 tuberculous meningitis patients from West Java using DNBSeq with 30x coverage. We excluded 3 individuals with inconsistent sex between genotype and phenotype data, 4 individuals visually identified as outliers in the principal component analysis (PCA) plot, and 5 individuals with high discordancy (>30%) between the WGS genotype count and the SNP-array genotype, leaving 227 samples for analysis. Within these genomes **(Supplementary Table 1)**, we identified 14,283,158 single nucleotide variants (SNVs) and 2,449,610 insertions and deletions (InDels). Among these, 6,616,414 (46.32%) SNVs and 941,524 (38.44%) InDels had a minor allele frequency (MAF) of less than 1%; 2,250,980 (15.76%) and 506,675 (20.68%) between 1% and 5%; and 5,415,764 (37.92%) and 1,001,411 (40.88%) more than 5%. All of the variants were then annotated using dbSNP build 153 as the reference. Based on the annotation, 1,867,419 (13.07%) SNVs and 432,345 (17.65%) InDels with MAF less than 1%; 22,122 (0.15%) and 200,711 (8.19%) with MAF between 1% and 5%; and 137 (0.001%) and 449,936 (18.37%) with MAF more than 5% were novel.

To understand the genomic region and function of the variants, we performed region-based and functional annotation using ANNOVAR. The majority of the variants was located in intronic and intergenic regions of the genome, consistent in all minor allele frequency (MAF) categories (**Figure 1**). The proportion of nonsynonymous SNV (nsSNV) among the total 14,283,158 SNVs, increased from 0.28% in the common (MAF > 5%) to 0.42 in the intermediate (MAF 1-5%, p <0001) and 0.63% in the rare (MAF < 1%) SNV categories (**Figure 1**).

**Figure 1.**
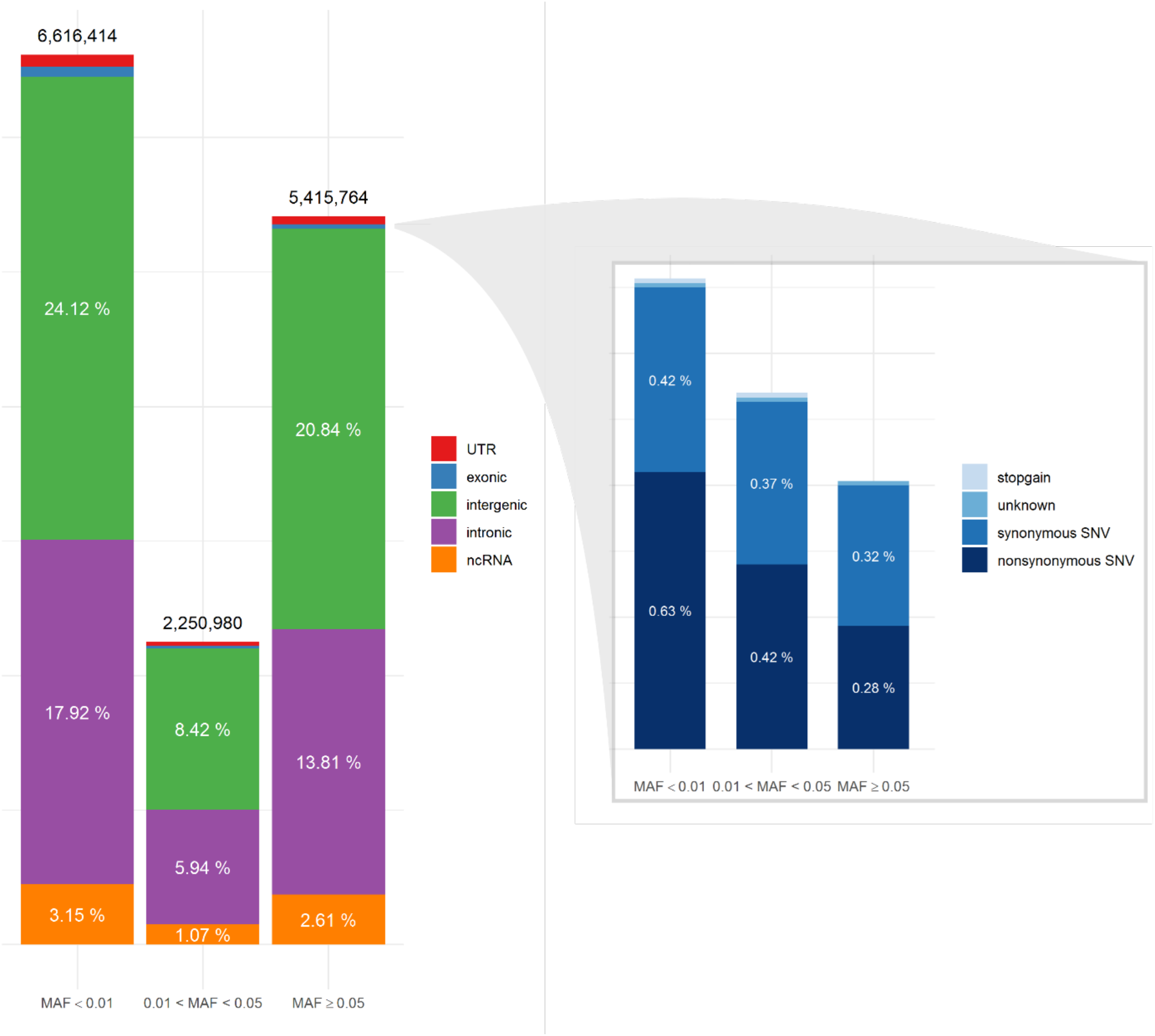
Single Nucleotide Variant (SNV) annotation utilizing ANNOVAR, displaying the distribution of SNV locations (*left*) and functional annotations of the exonic regions (*right*) for three minor allele frequency (MAF) categories.

### West Javanese population genetic architecture is distinct from other East Asian genomes

To evaluate the population structure of the West Javanese population, we performed Principal Component Analysis (PCA) including worldwide population reference data from the 1000 Genomes Project. The West Java population was found to be genetically closer to the Japanese, Chinese and Vietnamese individuals in the East Asian (EAS) cluster than to other 1000 Genomes populations, but did not completely overlap. **(Figure 2A)**. We also compared the genetic architecture of our West Javanese population to the individuals represented in the Genome Asia 100K (GAsP) project. These 68 individuals came from Flores, another island located approximately 1500 km east from West Java **(Figure 3)**, and comprises of four different ethnicities: Flores Bena, Flores Cibal, Flores Rampasana, and Austronesian. The West Javanese individuals were shown to be largely different from both the Austronesian population as well as the Flores Bena, Flores Cibal, and Flores Rampasana ethnicities which together form a third cluster (**Figure 2B**).

**Figure 2.**
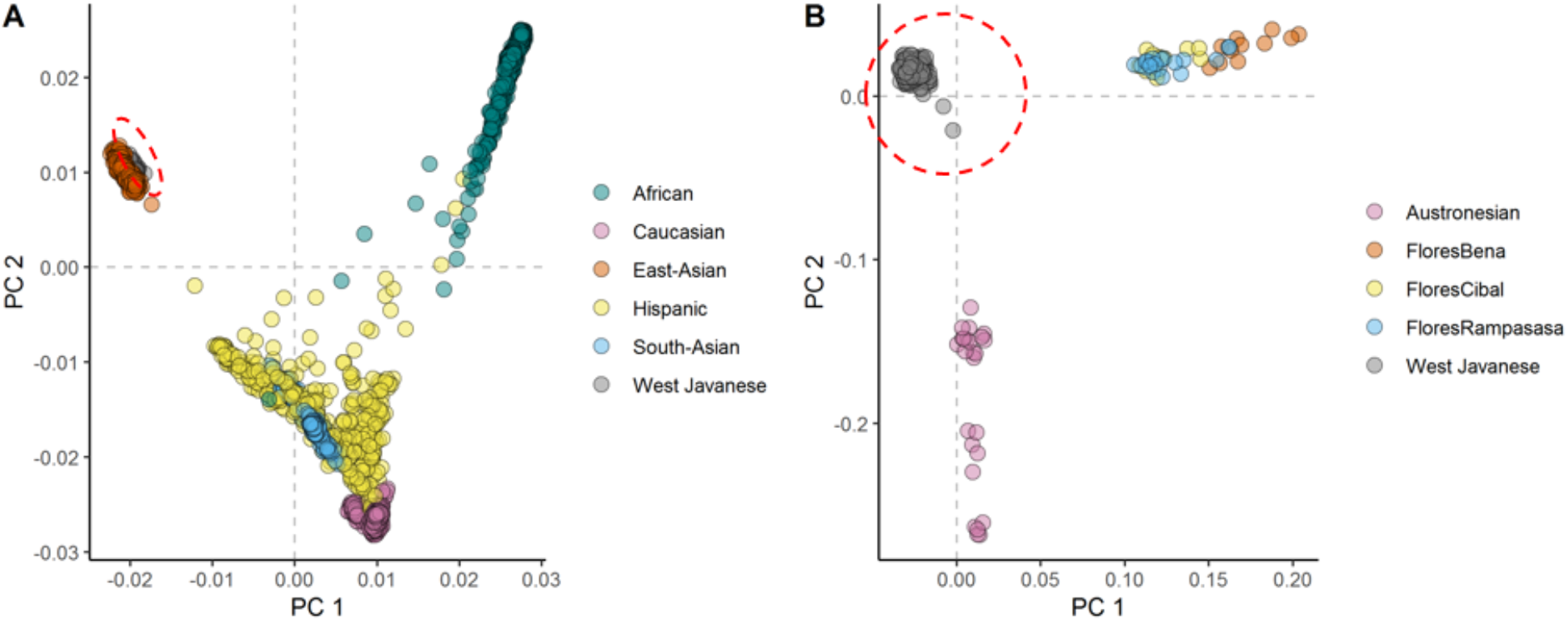
The first two principal components showing the genetic positioning of the West Javanese (red circle) population in relation to A) the 1000 Genome population and B) the Indonesian population (Flores) obtained from the *GenomeAsia* 100K project.

**Figure 3.**
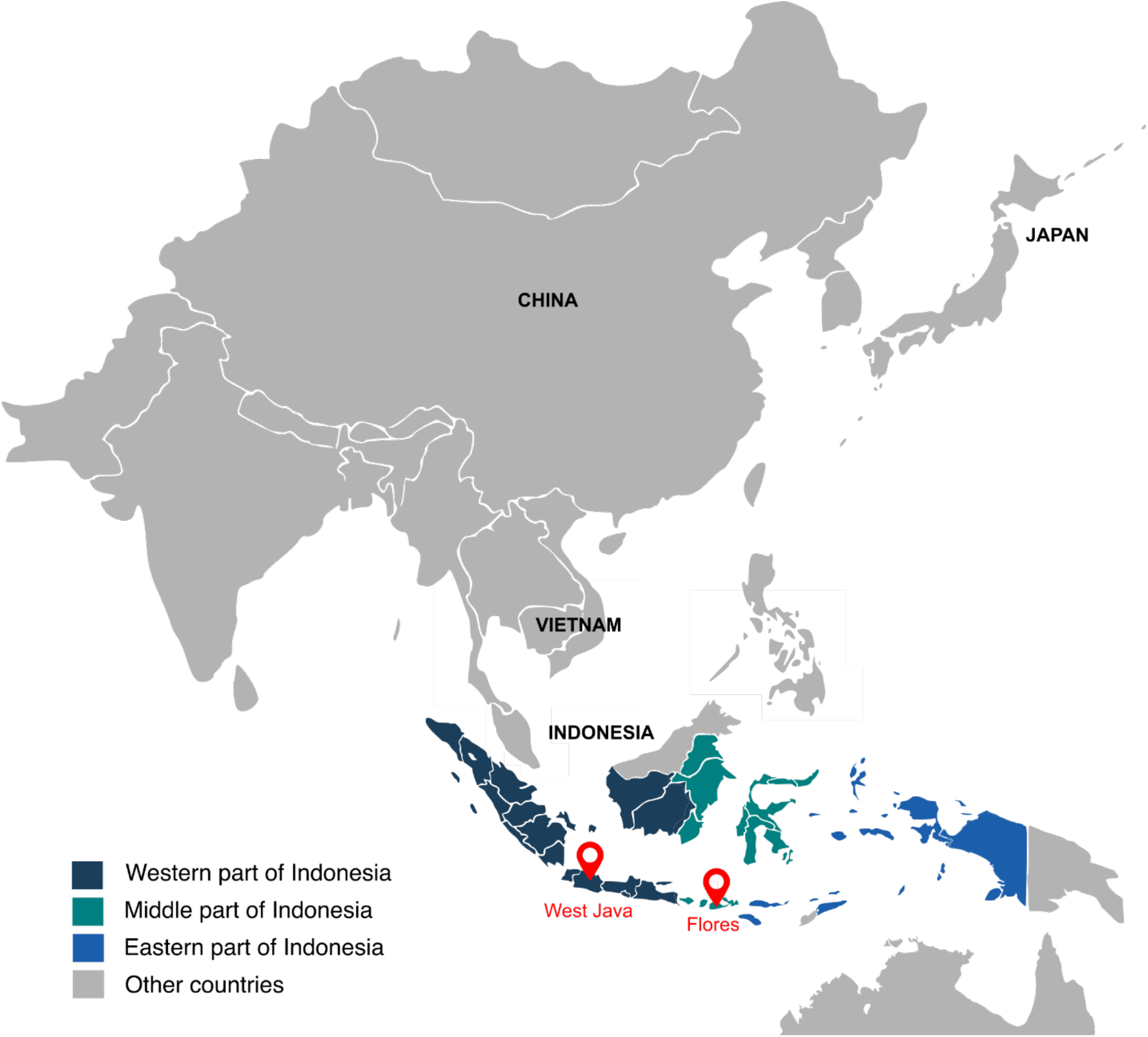
Map of Indonesia. This figure presents a map of Indonesia, highlighting the locations of West Java and Flores with two red marks. The East Asian panel from the 1000 Genome project includes individuals from China, Vietnam, and Japan.

To quantify the number of shared SNVs found in West Java population, with the Flores population and the populations in 1KGP3, we counted shared SNVs and categorized them by their MAF **(Table 1)**. Of the variants with a MAF < 0.01 in IND, less than half had been identified in both the Flores or the 1KGP3 populations. With increasing MAF, the proportion of known variants increased to approximately 90% (GAsP) and 98% (1KGP3). The larger representation in 1KGP3 pEAS could be explained because of a larger sample size, but also emphasizes the unexplored genetic diversity present within different ethnicities in Indonesia.

**Table 1.** Number of Single Nucleotide Variants (SNVs) identified in the genomes of West Javanese population, compared to the 100K Asia and 1000G populations.

| MAF | SNV (n) | Present in 100K Asia | Present in 1000G |
| --- | --- | --- | --- |
| < 0.01 | 6,818,363 | 1,093,308 (16.03%) | 2,975,954 (43.65%) |
| 0.01 - 0.05 | 2,263,232 | 1,445,656 (63.88%) | 1,962,389 (86.71%) |
| 0.05 - 0.1 | 1,054,398 | 922,880 (87.53%) | 1,023,669 (97.09%) |
| 0.1 - 0.2 | 1,393,726 | 1,248,343 (89.57%) | 1,358,766 (97.49%) |
| 0.2 - 0.3 | 1,109,694 | 999,800 (90.10%) | 1,088,726 (98.11%) |
| 0.3 - 0.4 | 966,199 | 869,030 (89.94%) | 950,003 (98.32%) |
| 0.4 - 0.5 | 922,765 | 829,212 (89.86%) | 908,100 (98.41%) |

### Comparison of imputed genotype and whole genome sequencing data reveal reduced accuracy

In addition to whole genome sequencing, 219 out of the 227 West Java individuals were also genotyped using a SNP-chip (HumanOmniExpressExome-8 v1.0; Illumina; San Diego, CA, USA). After imputation against the East Asian population in the 1KGP3 reference and QC check (R^2^ > 0.3 and MAF > 0.1), a final set of 4,751,257 SNVs was obtained.

To assess the accuracy of the imputed genotypes, treating our whole genome sequencing as the truth set, we compared the genotypes measured from the 2 different platforms for all 219 individuals. To visualize, we plotted the heterozygous allele count between the imputed SNVs and the whole genome sequencing **(Figure 4)**. As expected, SNVs with low R^2^ had larger discrepancies in heterozygous allele count between the imputed and whole genome sequencing. Importantly, even a cut-off of R^2^ greater than 0.8, leaves relevant discrepancies.

**Figure 4.**
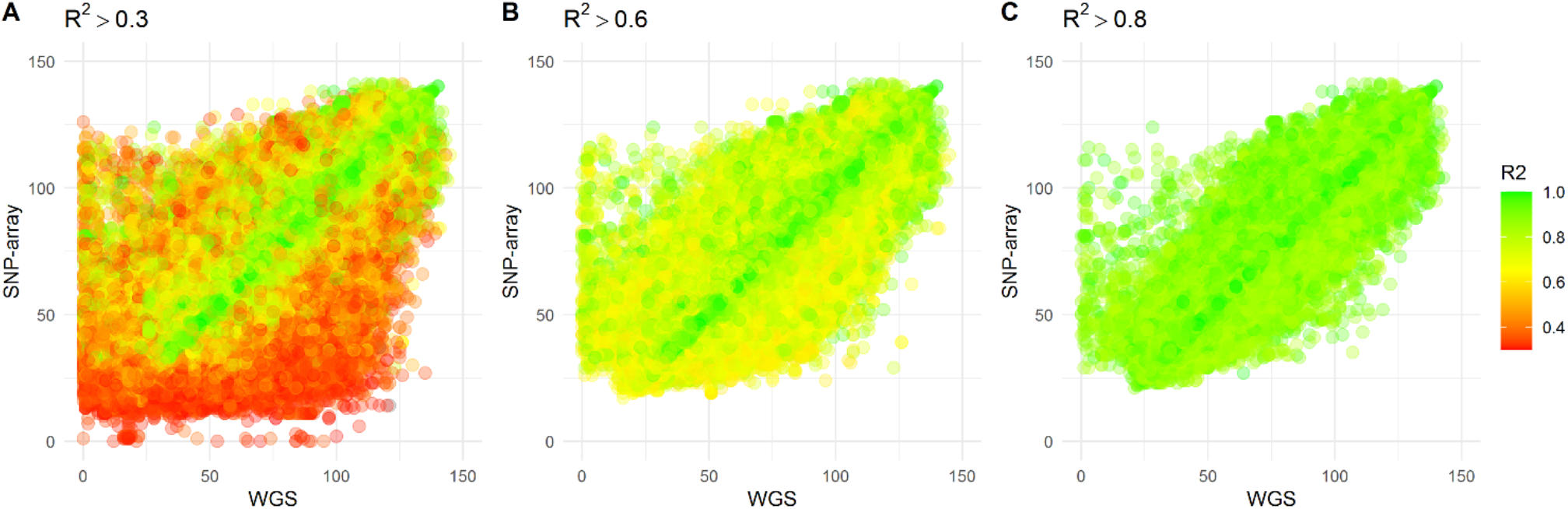
The scatter plots display the heterozygosity count of Single Nucleotide Variants (SNVs) from individuals who underwent both whole genome sequencing and SNP-array genotyping, imputed against the East Asian panel of the 1000 Genome project. Each dot on the plot represents an individual SNV, while the color indicates the imputation accuracy (R^2^) of the SNP-array data. The plots are presented based on three R2 thresholds: A) R^2^ > 0.3, B) R^2^ > 0.6, and C) R^2^ > 0.8.

### Imputation evaluation against EAS, IND, and EAS-IND panel

Inclusion of population-specific genome sequencing data into a reference panel has shown to increase the imputation accuracy ^10,11^. We aimed to assess the extent of accuracy improvement achievable by incorporating the West Java population into the existing 1KGP3 EAS panel. To accomplish this, we constructed three distinct reference panels: the 1KGP3 EAS panel (EASp), the Indonesian panel derived from West Java whole-genome sequencing (INDp), and the merged panel combining EAS and West Java datasets (EASp+INDp). Subsequently, we compared the imputation results of SNVs across all three reference panel configurations.

Out of 227 West Java WGS samples, 217 samples were utilized to create the reference panel consisting of INDp and EASp+INDp, and accuracy of imputation was evaluated for the three panels using two approaches: firstly, by comparing the R2 values (equivalent to the “info” metric in IMPUTE2). Ten remaining samples were used for benchmarking imputed SNVs against the WGS data by assessing the concordance between imputed SNVs and the WGS truth set in the 10 benchmark samples.

Imputation against EASp+INDp results in a higher count of SNVs with a total of 14,617,245 SNVs, compared to imputation to EASp (12,266,600 SNVs) or INDp (10,144,296 SNVs) individually. This outcome is in line with expectations, as merging two distinct panels increases the overall number of SNVs in the reference. Among the imputed SNVs, a subset of 7,792,202 SNVs is found in all three reference panels.

The EASp+INDp panel exhibited the highest mean R2 values, followed by the INDp and EASp panels **(Figure 5A)**, across the entire minor MAF spectrum. This finding suggests an enhanced imputation accuracy achieved by incorporating the Indonesian population. Subsequently, we assessed the genotype concordance between the imputed genotypes and the whole-genome sequencing (WGS) dataset using the 10 benchmarking samples. To ensure a precise evaluation of the concordance, we excluded genotypes that were expected to be imputed as homozygous reference. This step was crucial to prevent the misclassification of rare variants with low MAF as having high concordance, as the majority of the population is expected to have the homozygous reference genotype. We observed that the concordance of the imputed genotypes from the INDp and EASp+INDp panels were comparable, and both exhibited higher concordance rates compared to the EASp panel **(Figure 5B)**.

**Figure 5.**
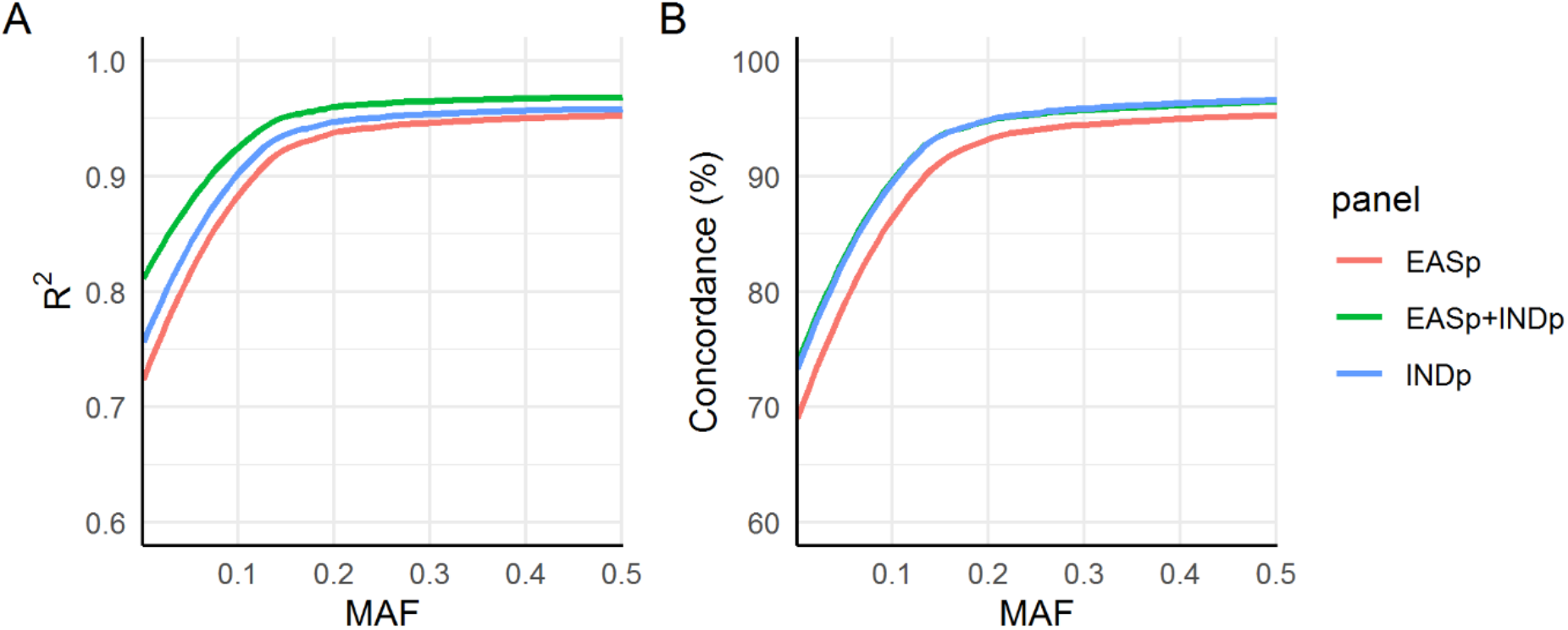
Imputation accuracy measurements. The figure presents a comparison of imputation accuracy across three panels using **A)** internal “INFO” metrics of IMPUTE2 and **B)** the actual genotype concordance between imputed single nucleotide variants (SNVs) and the whole-genome sequencing (WGS) dataset.

## Discussion

In this study, by examining the largest Indonesian cohort of genomes from individuals originating from West Java, we not only identified > 1.8 million novel variants but also highlight the rich genetic diversity present throughout the archipelago. Furthermore, by combining Indonesian and East Asian reference panels, we built an imputation reference panel to demonstrate significant improvement in SNP imputation accuracy and the concordance between imputed and genotyped variants. The implications of our findings are two-fold. Firstly, given the limited representation of non-European populations in genetic studies, combining the genetic architecture of this population can potentially help in understanding East-Asian population genetics. Additionally, the utilization of an imputation reference panel that incorporates specific populations becomes crucial to improve the imputation accuracy and in turn facilitate GWAS in diverse populations.

The high imputation accuracy observed among European GWA studies can be attributable to diverse representation of European populations in 1000 Genomes reference data. On the other hand, East Asian genomes from the 1000 Genomes data as well as the GenomeAsia data^4,12^ contains populations that are genetically closest to our study cohort. Although the GenomeAsia contains data from 219 population groups and 64 countries across Asia, our study clearly demonstrates that the Indonesian population possesses a distinctive genetic architecture compared to neighboring Asian countries. Furthermore, we identified a substantial number of novel rare variants that were absent in the 1000 Genomes East Asian population. These findings underscore the unique genetic profile of the Western Javanese population. Moreover, the results demonstrated a clear distinction between the genetic makeup of the Western Javanese population and populations originating from the central region of Indonesia.

It is not surprising that adding the Western Javanese population WGS data to the reference panel improved the imputation accuracy of common variants in our study. In fact, several previous studies have already shown the positive impact on imputation performance when additional populations were combined in reference panels^13–15^. However, this improvement in accuracy also varied depending on the allele frequencies of the variants, where much stronger improvement was seen for common variants (MAF > 0.1). It is shown that in addition to the use of large number of samples to build reference panels^16,17^, using population-specific reference panels^18,19^ strongly benefit rare variant imputation. As GenomeAsia currently has a small sample size of 68 individuals from Indonesia, our study helps to enrich the Asian reference datasets further to assist in rare variant imputation.

A notable strength of our study lies in the substantial sample size employed to construct the reference panel, consisting of 217 individuals. The inclusion of a large number of individuals allows for a more comprehensive representation of haplotypes within the population, consequently improving the accuracy of imputation. A potential limitation of our study is that the population in our study comprises of patients diagnosed with tuberculous meningitis, and this selection may impact the distribution and representation of SNPs within the population. This may especially be relevant for rare variants that would put individuals at risk for tuberculous meningitis but less so for variants that are highly polymorphic (MAF>0.1). Given that the number of disease-affecting loci is small compared to the entire genome, capturing population-specific aspects should not be significantly impacted^20^. As cost of WGS is reducing, future studies should ideally include healthy individuals to mitigate the potential selection bias effects and further validate our findings.

In conclusion, our study provides evidence that the genetic architecture of the Indonesian population exhibits distinct characteristics when compared to other Asian countries. Our study also serves as an important resource to improve the GWAS of complex phenotypes in the West Javanese population. Given the rich ethnic diversity in the country, it is crucial for future genetic research to encompass a wider range of Indonesian ethnicities to capture the full extent of genetic diversity present in the country. By expanding our knowledge of the genetic architecture within Indonesia, we can pave the way for more targeted and effective precision medicine strategies tailored to the diverse needs of different ethnic groups.

## Supporting information

Number of rare (MAF < 0.01) Single Nucleotide Variants (SNVs) and Insertions/Deletions (InDels) in the West Javanese Whole Genome Sequencing dataset

## Acknowledgements

The authors thank Alifah Taqiya, Fransisca Kristina, and Rani Trisnawati for coordinating sample and data management; the director of the Hasan Sadikin General Hospital, Bandung, Indonesia, for accommodating the research. We also express our gratitude to our funder: This study was supported by National Institutes of Health (R01AI145781). The funder had no role in study design, data collection and analysis, decision to publish, or preparation of the manuscript.

## Author contributions

EA, ALR, AvL, RvC, and VK conceptualized the study design. EA performed the analysis. SD, ARG, BA, TPS contributed to manuscript preparation. All authors reviewed and approved of the manuscript.

## Supplemental information

Table S1

## STAR Methods

### Whole genome sequencing and data processing

As part of the ULTIMATE project^21^, we sequenced DNA from 239 tuberculous meningitis patients. The genomic DNA sample was fragmented randomly by Covaris technology, resulting in fragments of 350bp after selecting the appropriate size range. The fragmented DNA was subjected to end repair, followed by the addition of an “A” base at the 3’-end of each strand. Adapters were then ligated to both ends of the DNA fragments and subjected to amplification by ligation-mediated PCR (LM-PCR), followed by single-strand separation and cyclization. Rolling circle amplification (RCA) was used to generate DNA Nanoballs (DNBs) from the qualified DNBs, which were loaded onto patterned nanoarrays and pair-end reads were obtained on the DNBseq platform. High-throughput sequencing was performed for each library to ensure adequate sequencing coverage. The raw image files generated during sequencing were processed by DNBseq base calling software using default parameters to obtain sequence data in the form of paired-end reads, which is defined as “raw data” and stored in FASTQ format.

FASTQ files of each sample was first mapped to the human reference genome (NCBI Build 37, hg19) (*https://www.ncbi.nlm.nih.gov/assembly/GCF_000001405.13/*) using BurrowsWheeler Aligner (BWA)^22^. To ensure reliable results, we followed the recommended Best Practices for variant analysis with the Genome Analysis Toolkit (GATK)^23–25^. Local realignment around InDels and base quality score recalibration was performed using GATK v3.7-0^23–25^, with duplicate reads removed by Picard tools^26^. GVCF files for each sample were then created using HaplotypeCaller. All the individual GVCFs were then jointly genotyped using GATK GenotypeGVCFs, after first combining them with GATK CombineGVCFs. Variants were further recalibrated using GATK Variant Quality Score Recalibration (VQSR), and variants not passing the VQSR filtering and have missingness > 5% were removed. Region-based and functional annotation were done using ANNOVAR version 7 June 2020^27^.

### Whole genome sequencing data of Genome Asia 100K

After receiving approval from the 100K GenomeAsia consortium^9^, VCF files of the Indonesian population were obtained, consisting of 68 individuals. In the final VCF files containing variant sites for Indonesian population, non-variant sites with an allele count of 0 were excluded.

### Imputation reference panel creation

Using IMPUTE2^28^, we created three imputation reference panels: The East Asian panel (EASp), Indonesian panel (INDp), and the combined East Asian and Indonesian panel (EASp+INDp). Out of 227 Indonesian whole genome sequences (WGS) that passed QC, 10 Indonesian WGS were taken out for benchmarking purpose, and the remaining 217 were used to create the reference panel. To obtain the EASp, the publicly available reference panel The 1000 Genomes phase 3 (1KGP3)^4^ was downloaded and the East Asian population was extracted. For both EAS and IND WGS data, multiallelic sites were split into biallelic sites, and variants with missingness > 5%, Hardy Weinberg Equilibrium (HwE) < 1x10^−10^, and allele count < 2 were removed. The 217 IND WGS were then phased using SHAPEITv2^29^ and converted to a reference panel using IMPUTE2^28^. The EAS and IND panel (EASp+INDp) was constructed by merging the EAS and IND panel using the “-merge_ref_panels_output_ref” option in IMPUTE2^28^, which performed the merging in two steps: first, imputing variants that were specific to one panel to the other panel and vice versa, and second, combining the two panels by taking the union of variants from both panels.

### Imputation against IMPUTE2 reference panel

Genotypes of 509 TBM patients established with SNP-typing (HumanOmniExpressExome-8 v1.0; Illumina; San Diego, CA, USA) were used as data input for imputation. Variants with >5% missingness or HwE < 0.00001 were removed, and then pre-phased using SHAPEITv2^29^. To align the strand against each of the reference panel, we used GenotypeHarmonizer^30^, which exploit linkage disequilibrium pattern to solves the unknown strand issue by aligning ambiguous A/T and G/C SNPs to a specified reference, thus eliminating the need of prior knowledge of the used strand.

Imputation was performed against the 3 panels using IMPUTE2^28^ by first dividing the input genome into 5Mb chunks to increase computation and memory efficiencies. After completion of imputation, all chunks were re-combined to obtain the final imputed genotype, in IMPUTE2 haplotype format. The haplotype files were converted to VCF for further analysis using SHAPEITv2^29^.

### Evaluation of Imputation Accuracy

To assess the imputation accuracy, we first utilized the internal quality metrics obtained from IMPUTE2, specifically the INFO score. Subsequently, we calculated the genotype concordance of the overlapping single nucleotide variants (SNVs) between the imputed SNVs and the whole-genome sequencing (WGS) dataset using *vcfcompare* tools. For an accurate evaluation of concordance, we excluded SNVs expected to be homozygous reference and calculated the concordance solely based on the alternate allele.

