## Supplementary material for "Sequencing whole genomes of the West Javanese population in Indonesia reveals novel variants and improves imputation accuracy": Number of rare (MAF < 0.01) Single Nucleotide Variants (SNVs) and Insertions/Deletions (InDels) in the West Javanese Whole Genome Sequencing dataset: Supplementary Table 1.pdf

**Supplementary Table 1** Number of rare (MAF < 0.01) Single Nucleotide Variants (SNVs) and Insertions/Deletions (InDels) in the West Javanese Whole Genome Sequencing dataset. The number and percentage of variants detected by WGS are indicated.

| chr | SNV | | | InDel | | |
|---|---|---|---|---|---|---|
| | n | MAF<0.01 (%) | Novel (%) | n | MAF<0.01 (%) | Novel (%) |
| 1 | 1,142,385 | 542,341 (47.47) | 159,050 (13.92) | 203,229 | 77,109 (37.94) | 91,251 (44.90) |
| 2 | 1,125,085 | 493,890 (43.90) | 131,482 (11.69) | 217,566 | 86,852 (39.92) | 96,341 (44.28) |
| 3 | 998,139 | 447,438 (44.83) | 125,241 (12.55) | 175,037 | 68,860 (39.34) | 77,211 (44.11) |
| 4 | 1,029,070 | 463,698 (45.06) | 132,752 (12.90) | 171,690 | 66,233 (38.58) | 73,229 (42.65) |
| 5 | 912,383 | 418,083 (45.82) | 121,971 (13.37) | 156,121 | 62,126 (39.79) | 68,222 (43.70) |
| 6 | 923,747 | 442,157 (47.87) | 132,683 (14.36) | 149,907 | 57,031 (38.04) | 64,322 (42.91) |
| 7 | 811,295 | 359,619 (44.33) | 103,287 (12.73) | 148,801 | 58,230 (39.13) | 65,590 (44.08) |
| 8 | 832,534 | 407,651 (48.97) | 124,267 (14.93) | 124,224 | 49,907 (40.18) | 54,998 (44.27) |
| 9 | 641,048 | 298,000 (46.49) | 88,948 (13.88) | 101,379 | 38,170 (37.65) | 44,857 (44.25) |
| 10 | 713,173 | 322,913 (45.28) | 88,164 (12.36) | 118,280 | 44,339 (37.49) | 52,154 (44.09) |
| 11 | 697,223 | 324,751 (46.58) | 90,386 (12.96) | 113,123 | 43,917 (38.82) | 49,546 (43.80) |
| 12 | 667,314 | 298,554 (44.74) | 79,976 (11.98) | 123,633 | 47,061 (38.07) | 55,351 (44.77) |
| 13 | 528,819 | 249,104 (47.11) | 71,526 (13.53) | 89,867 | 34,850 (38.78) | 38,011 (42.30) |
| 14 | 479,730 | 225,262 (46.96) | 64,511 (13.45) | 82,810 | 31,675 (38.25) | 36,192 (43.70) |
| 15 | 459,534 | 227,984 (49.61) | 68,859 (14.98) | 75,693 | 29,030 (38.35) | 33,711 (44.54) |
| 16 | 446,985 | 211,452 (47.31) | 57,336 (12.83) | 69,332 | 25,029 (36.10) | 32,502 (46.88) |
| 17 | 411,226 | 202,703 (49.29) | 58,750 (14.29) | 78,585 | 29,261 (37.23) | 37,932 (48.27) |
| 18 | 404,206 | 185,227 (45.82) | 53,341 (13.20) | 64,926 | 24,300 (37.43) | 27,625 (42.55) |
| 19 | 337,343 | 155,476 (46.09) | 41,244 (12.23) | 63,924 | 21,385 (33.45) | 30,532 (47.76) |
| 20 | 315,433 | 149,288 (47.33) | 43,024 (13.64) | 53,219 | 20,202 (37.96) | 24,049 (45.19) |
| 21 | 203,343 | 94,607 (46.53) | 26,070 (12.82) | 33,382 | 12,999 (38.94) | 13,629 (40.83) |
| 22 | 203,143 | 96,216 (47.36) | 26,810 (13.20) | 34,882 | 12,958 (37.15) | 15,737 (45.11) |
| **Total** | **14,283,158** | **6,616,414 (46.32)** | **1,889,678 (13.23)** | **2,449,610** | **941,524 (38.44)** | **1,082,992 (44.21)** |
